# HIV corruption of the Arp2/3-Cdc42-IQGAP1 axis to hijack cortical F-Actin to promote cell-cell viral spread

**DOI:** 10.1101/2019.12.13.873612

**Authors:** Anupriya Aggarwal, Alberto Ospina Stella, Catherine Henry, Kedar Narayan, Stuart G. Turville

**Affiliations:** The Kirby Institute, University of New South Wales, New South Wales, Australia; Center for Molecular Microscopy, Center for Cancer Research, National Cancer Institute, National Institutes of Health, Bethesda, Maryland, USA; Cancer Research Technology Program, Frederick National Laboratory for Cancer Research, Frederick, Maryland, USA

## Abstract

F-Actin remodelling is important for the spread of HIV via cell-cell contacts, yet the mechanisms by which HIV corrupts the actin cytoskeleton are poorly understood. Through live cell imaging and focused ion beam scanning electron microscopy (FIB-SEM), we observed F-Actin structures that exhibit strong positive curvature to be enriched for HIV buds. Virion proteomics, gene silencing, and viral mutagenesis supported a Cdc42-IQGAP1-Arp2/3 pathway as the primary intersection of HIV budding, membrane curvature and F-Actin regulation. Whilst HIV egress activated the Cdc42-Arp2/3 filopodial pathway, this came at the expense of cell-free viral release. Importantly, release could be rescued by cell-cell contact, provided Cdc42 and IQGAP1 were present. From these observations we conclude that a proportion out-going HIV has corrupted a central F-Actin node that enables initial coupling of HIV buds to cortical F-Actin to place HIV at the leading cell edge. Whilst this initially prevents particle release, maturation of cell-cell contacts signals back to this F-Actin node to enable viral release & subsequent infection of the contacting cell.

## Introduction

Actin is a major component of the cellular cytoskeleton and is present in both monomeric globular (G-actin) and polymeric filamentous (F-actin) forms in all eukaryotic cells. Specifically in human leukocytes, actin accounts for over 10% of the total protein content and is a prerequisite for many pathways involved in communication of the immune response, such as chemotaxis of leukocytes through to the formation of supramolecular structures like the immunological synapse-a fundamental structure driving the primary immune response. While cells encode a wide range of proteins that mediate F-actin remodelling, the critical ability to seed or ‘nucleate’ F-actin from the monomeric G-actin pool is limited to only a few protein families. The two major classes of cellular actin nucleators are the Arp2/3 complex and formins. The Arp2/3 complex is composed of 7 different subunits and allows formation of branched actin networks, through nucleation of a branch filament from an existing mother filament at an angle of 70 degrees [1]. In contrast, formins are large multidomain proteins that drive nucleation and/or elongation of unbranched linear actin filaments. The activity of cellular Arp2/3 and formins is tightly regulated by a complex network of signalling pathways that primarily rely on the molecular switch properties of Rho-GTPases, such as Rac1 and Cdc42, for their activation [2].

HIV infection and spread proceeds primarily in CD4 positive leukocytes of our immune system. Viral spread can be observed at two levels. Firstly, free virus release from infected cells, with virions travelling in a cell-free form until encountering a new target cell to infect. Secondly, HIV budding that occurs directly at sites of cell-cell contact. The supramolecular structure that enables the latter and highly efficient cell-cell viral transfer is referred to as the virological synapse (VS) [3,4]. In both cases, viral budding needs to proceed at the plasma membrane (PM) of infected cells and is initially driven by oligomerisation of the HIV structural protein Gag [5] and culminates with HIV particle abscission mediated by cellular proteins of the endosomal sorting complexes required for transport machinery (ESCRT) [6]. Several F-actin structures have been previously observed in association with HIV assembly and higher-level Gag oligormerisation. These include; the temporal formation of F-Actin asters/stars that appear just underneath the PM prior to particle release [7], and assembling HIV particles decorating the tips of finger-like filopodial structures [8-10]. It is however unclear how these events are mechanistically connected and how coupling to the F-actin cytoskeleton benefits HIV release/spread. This can be considered at two-levels: firstly how does F-Actin regulation influence the assembly and release of cell free virus in infected cells? For instance, do cortical F-Actin structures facilitate HIV assembly and release at the PM as observed for other viruses [11]? Secondly, how is F-actin regulated at cell-cell contacts involving HIV infected cells? Several studies have shown that functional actin dynamics are required for cell-cell viral transfer [4], yet how HIV assembly and release are spatiotemporally coordinated during this process has not been clarified mechanistically.

With our primary aim to determine the role for F-Actin in cell-free HIV egress and cell-cell viral transfer, herein we peeled back the complexity of F-Actin regulation in leukocytes by successive depletion and/or knockout of key actin nucleators and other associated proteins that regulate their activity. In doing so, we biased the formation of different cortical F-Actin structures such as filopodia and lamellipodia. Herein we define these structures as outlined by Mattila and Lappalain, i.e. Filopodia are cylindrical finger like protrusions approximately 100–300 nm in diameter and up to 1μm to 10 μm or more in length, whereas lamellipodia are thin (100 nm to 200 nm thick) sheet/viel like cortical F-Actin protusions [12]. Whilst the regulation of lamellipodia is well understood and primarily depends on branched F-actin nucleation by the Arp2/3 complex downstream of Rac1 and its effector Wave2 [13,14], various models have been proposed for filopodia formation and increasing evidence suggests this process may be cell-type specific [15]. Importantly, little is known about the mechanism of filopodia formation in cells of hematopoietic lineage, despite the fact that filopodia play important roles in immune cell function.

Using a combination of live cell imaging, focused ion beam scanning electron microscopy (FIB-SEM), virion mass spectrometry and viral infection assays, we observed the influence of HIV on cortical F-Actin at several levels. First, FIB-SEM revealed HIV budding to be relatively enriched in areas of high positive membrane curvature within Arp2/3-dependent cortical F-Actin structures, including filopodia and lamellipodia. Second, virion mass spectrometry identified a cortical F-Actin signalling node comprising of the Arp2/3 related GTPases Rac1 and Cdc42 and their binding partner, the scaffolding protein IQGAP1. Finally, while depletion of a number of dominant F-actin regulators was observed to affect free virus release, cell-cell viral transfer was only significantly impaired in cells depleted of Cdc42 or IQGAP1. Collectively these observations support a dominant role of the GTPase Cdc42 and IQGAP1 in the final stages of viral egress and cell-cell spread. In this setting we propose HIV manipulation of the Cdc42/IQGAP1 node to be important at two levels: firstly it enables HIV to be embedded and retained in Arp2/3-dependent leading edge structures that are important during pre-synaptic events. Secondly, as the VS is engaged and matures, the same regulators likely coordinate F-Actin dynamics to enable conditions that facilitate final viral particle release.

## Results

### Moulding cortical F-Actin through formin and Arp2/3 depletion

A physical association of HIV with F-actin structures has been previously observed in all major HIV primary target cell types [9,16]. In infected CD4+ T-cells and dendritic cells this manifests in the form of HIV-Filopodia, which are F-actin rich finger-like structures with HIV assembly observed at their tips [9]. Since these structures are more prominent on dendritic cells and similarly enriched in U937 cells [9,17,18], this latter myeloid cell line provides an ideal model to dissect the link between F-Actin and HIV assembly in the specific context of hematopoietic cell lineages. Given the proposed role of Arp2/3 and formins in filopodia formation in other cell types, we initially focused on these key actin regulators for depletion. However, while Arp2/3 is ubiquitously expressed in eukaryotic cells, there are at least 15 different formins in vertebrates [19]. Since Diaph1, Diaph2 and FMNL1 are the most abundantly expressed in leukocytes, we tailored our initial shRNA screening to depletion of these formins. Disruption of filopodial networks was assessed by measuring filopodial abundance (average number of filopodia per cell) and length (average filopodial length measured from the PM to the tip). To this end we used several imaging techniques at increasing levels of resolution, including; i) live cell imaging, ii) fixed cell fluorescence imaging followed by 3D deconvolution, and iii) the power of correlative FIB-SEM to finely resolve F-Actin structures closer to the PM. In brief, FIB-SEM represents a method where iterative cycles of finely tuned ion abrasion milling are followed by high-resolution scanning electron microscopy of heavy-metal stained, resin-embedded cell samples [20,21]. The end result is the recording of a stack of 2D back-scatter electron images, which are then processed and converted to a 3D image volume, typically at ∼ 10 nm pixel sampling (Fig.1 B-E). This method provides a powerful imaging tool for cell biology and virology, as it gives users the ability to resolve nanoscale ultrastructural features in cellular samples that may appear in association with viral particles [22].

**Figure 1.**
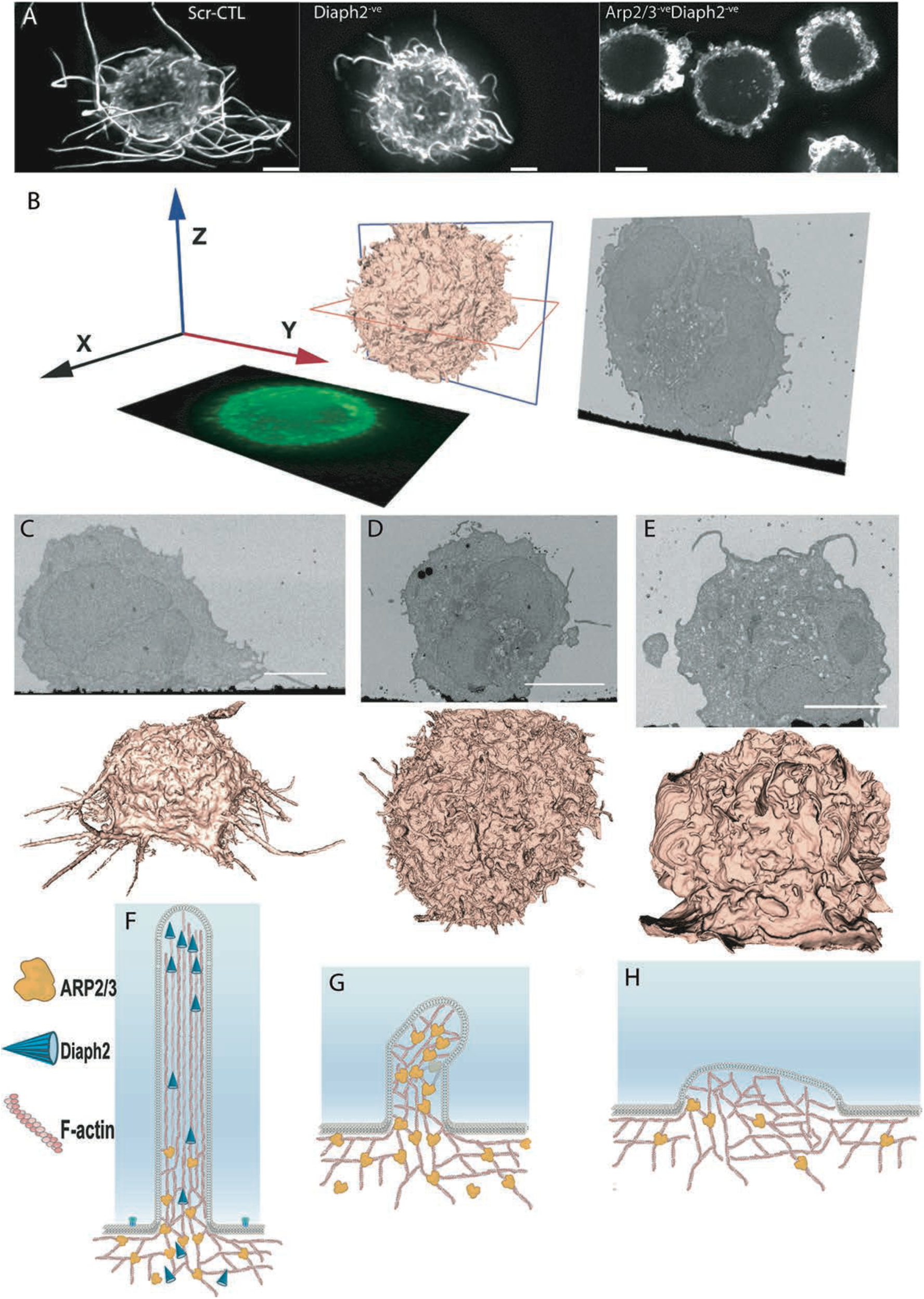
Depletion of F-actin regulators reveals long filopodial networks to be driven by Arp2/3 and Diaph2. A. F-Actin staining by phalloidin Alexa-647 reveals extensive and long filopodia in uninfected scramble shRNA control cells (Scr-CTL), short abundant filopodia upon deletion of the formin Diaph2, and extensive lamellipodia when both Diaph2 and Arp2/3 are co-depleted. B. Schematic representation of FIB-SEM imaging data. C-E. FIB-SEM 2D images (upper) and 3D reconstructions (lower) of C. non-depleted control cells (Scr-CTL), D. Diaph2-depleted cells and E. Diaph2 and Arp2/3 co-depleted cells. All scale bars are 5μm. F-H. Schematic representation of phenotypes induced by loss of F-Actin regulators. F. Wildtype scenario, G. Diaph2 deficiency, H. Diaph2 & Arp2/3 deficiency. Note in H. we represent reduced levels of Arp2/3, given its high cellular abundance and residual levels of Arp2/3 in our knockdown cells (Fig. S3).

Initial experiments revealed filopodial lengths to be dependent solely on the formin Diaph2 and not the other leukocyte-enriched formins (Fig. S1). Silencing of Diaph2 by shRNA achieved >95% depletion at the protein level (Fig. S3), and in these cells cortical F-Actin coalesced into a network of abundant and short (1 to 3µm) filopodia (Fig. 1E&H; Movie S1). Since we could confirm this phenotype in CRISPR/Cas9-generated and clonally expanded Diaph2 homozygous knockout cells, our observations suggest that filopodial length but not seeding is dependent on Diaph2. Subsequent shRNA codepletion of other expressed formins in addition to Diaph2 did also not disrupt this short filopodial network (Fig. S1). Therefore, to test if the shorter but more abundant filopodia were Arp2/3-dependent, we disrupted both Diaph2 and the Arp2/3 complex by shRNA. Co-depletion led to short filopodia converging into an extensive lamellipodial network (Fig. 1A,E&H; Movie S2). To conclude, we could readily control cortical F-Actin within this leukocyte landscape, and generate three unique cell types with a continuum of cortical F-actin structures; i) long filopodia, ii) abundant short filopodia, and iii) an extensive network of large lamellipodia. Furthermore, our observations indicate that seeding of filopodia in myeloid cells requires Arp2/3-mediated actin nucleation, whereas filopodial elongation is dependent on the formin Diaph2 (Fig. 1. F-H).

### The influence of shifting F-Actin structures on the location of HIV budding

In the context of HIV infected cells, we used our high-resolution imaging approaches to probe for a possible link between HIV assembly and specific F-Actin structures and/or pathways in leukocytes. Previously we have observed live cells with long filopodia to have significant numbers of HIV positive tips [9], and readily concluded that in untreated cells HIV assembly is enriched to this site. However, detection of HIV buds in cells with short filopodia (i.e. depleted of Diaph2) was constrained by the inability to resolve F-Actin structures proximal to the PM by fluoresecence microscopy alone. Thus, we applied FIB-SEM imaging to HIV-infected Diaph2^-ve^ cells (short filopodia) and observed HIV buds in routine association with the tip and sides of these structures (Fig. 2A,B,E&H). In cells with prominent lamellipodia (Diaph2^-ve^Arp2/3^-ve^), FIB-SEM imaging revealed abundance of HIV buds along the ridges of lamellipodia (Fig. 2F&K; Movie S2). Therefore one common feature of each F-Actin structure was the appearance of HIV preferentially in areas of positive membrane curvature. Given the large topological differences between filopodia and lamellipodia, we assessed viral-bud density by accounting for the surface area available for budding within each distinct F-Actin structure. This revealed HIV buds to be significantly enriched in areas of high positive membrane curvature (Table SI; Fig. 2 G-L): Lamellipodial ridges and filopodia were the most active areas of viral assembly, with a distinct preference for the latter (filopodia tips outscored lamellipodial ridges by 5-fold, when surface area was considered). This observation supports two potential mechanisms. First, HIV assembly is facilitated by areas of positive curvature or alternatively, HIV assembly recruits/influences cellular protein(s) at the PM that can lead to positive curvature.

**Figure 2.**
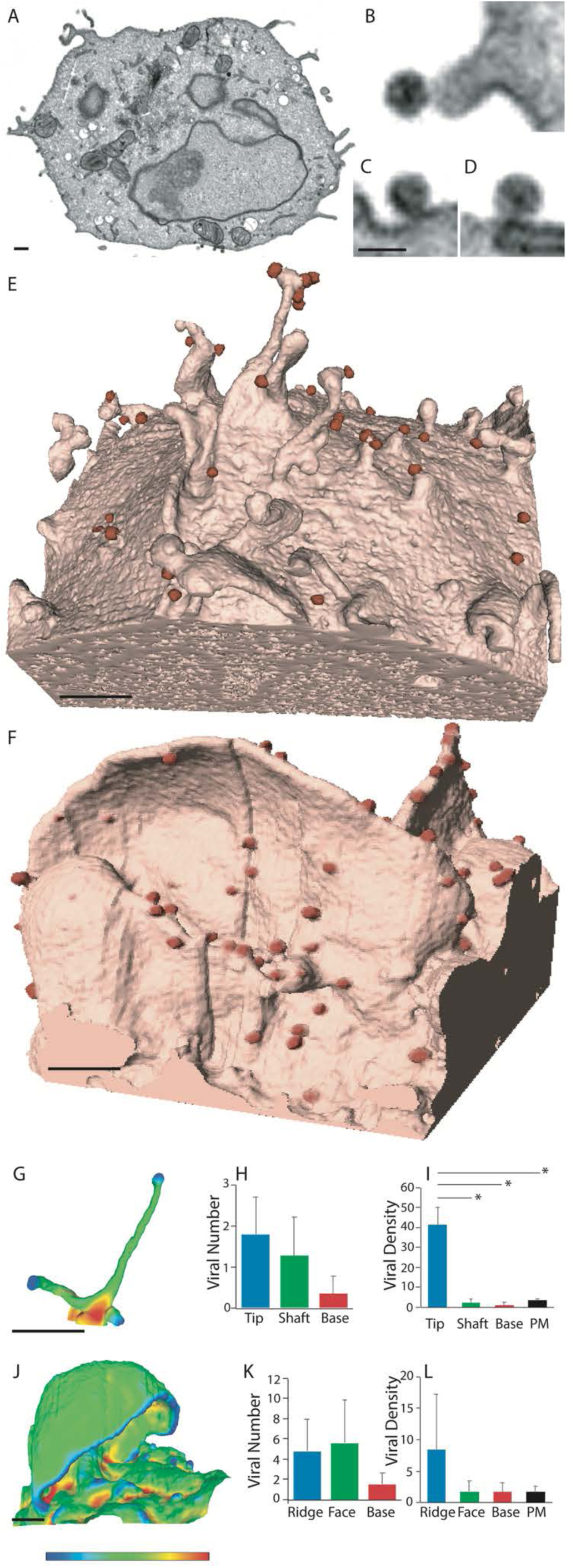
HIV budding enriched to positively curved cortical F-Actin. A-D Representative FIB-SEM images of HIV virions associated with B. Filopodia and C-D plasma membrane of a HIV infected cell depleted of Diaph2. E.-F. 3D rendering FIB-SEM images of HIV infected E. Diaph2^-ve^ and F. Diaph2^-ve^Arp2/3^-ve^ cell clones. HIV buds are shaded in red to highlight their location. G&J. enumeration of HIV-buds in association with positively curved cortical F-Actin structures. G. Filopodia and J. lamellipodia are pseudocolored using a colour spectrum from blue (positive curvature), green (neutral curvature) to red (negative curvature). H-I Enumeration of total HIV buds in association with the filopodia in Diaph2^-ve^ cells. H. Absolute viral bud counts and I. HIV bud density per μm^2^. K-L. Enumeration of total HIV buds in association with the lamellipodia in Diaph2^-ve^ Arp2/3^-ve^ cells. K. Absolute viral bud counts and L. HIV bud density per μm^2^. All scale bars are 0.5μm with the exception of B-D, which is 0.1μm. Bar graphs in H-I represent mean and standard deviations of virion counts from n = 15 filopodial structures. K-L bar graphs represent mean and standard deviations of virion counts from n = 15 lamellipodial structures. Statistics for panels G-L are summarised in Table SI. *P>0.05

### Filopodia dominated by the formin Diaph2 present positive curvature at the plasma membrane but exclude assembling HIV particles

To assess whether strong positive membrane curvature alone was sufficient to position HIV buds at filopodial tips, we induced long filopodia using a constitutively active (C/A) mutant of Diaph2. Diaphanous-related formins exist in an autoinhibited conformation mediated by the interaction between their N-proximal inhibitory domain (DID) and C-terminal autoregulatory domain (DAD) [23]. Disruption of this autoinhibitory state can be achieved by formins binding to Rho-GTPases or, as in our case, by deletion of their C-terminal DAD domain [24]. Importantly, in both cases the central actin polymerization domain of the formin is rendered constitutively active.

If HIV assembly was directly promoted by positive membrane curvature, filopodia induced by Diaph2^C/A^ would incorporate assembling viral particles. However, while Diaph2^C/A^ expression readily induced the formation of long straight filopodia with Diaph2 accumulating at the filopodial tips (Fig. 3A & Movie S3), in HIV infected cells we also observed complete exclusion of HIV particles from the tips of these structures (Fig. 3A-C & Movie S3). Therefore, the strong membrane curvature in filopodial tips alone is not sufficient to recruit HIV assembly to this region. Since long filopodia in WT cells are routinely HIV positive, whereas straight C/A Diaph2 driven filopodia are not, it is unlikely that formins represent the link of HIV to the F-actin cytoskeleton. We then turned our attention to the Arp2/3 complex, given that previous observations propose this as the dominant F-actin nucleator at the cell cortex, with formin activity being restricted to filament elongation post F-actin nucleation [25]. To confirm if Arp2/3 was the dominant filopodial nucleator in WT versus Diaph2^C/A^ cells, we immunostained filopodia for Arp2/3, and examined the footprint of this nucleator along the filopodial body and tip. In both cell types, the filopodial bases (3μm from the membrane) were all Arp2/3 positive (Fig. 3D & E). In contrast, the filopodial tips of Diaph2^C/A^ cells were negative for Arp2/3 antigen (Fig. 3D), whereas Arp2/3 was frequently observed along the entire shaft and at the tip of wildtype filopodia (Fig. 3E). To quantify the extent of Arp2/3 tip exclusion, we measured the distance from the tip of filopodia to the first detectable Arp2/3 signal and observed a significantly greater distance of Arp2/3 from the filopodial tip in Diaph2^C/A^ cells relative to WT cells (6.2μm versus 1.4μm; *p*>0.0001; *n* = 50). In summary, by mapping the HIV budding sites at high resolution we could reach several conclusions. Firstly, HIV buds primarily enrich to cortical F-Actin structures with positive curvature. Secondly, positive curvature and/or Diaph2 activity alone are not responsible for the enrichment of HIV buds to these sites. Finally, Arp2/3-dependent cortical F-actin structures are primarily HIV positive.

**Figure 3.**
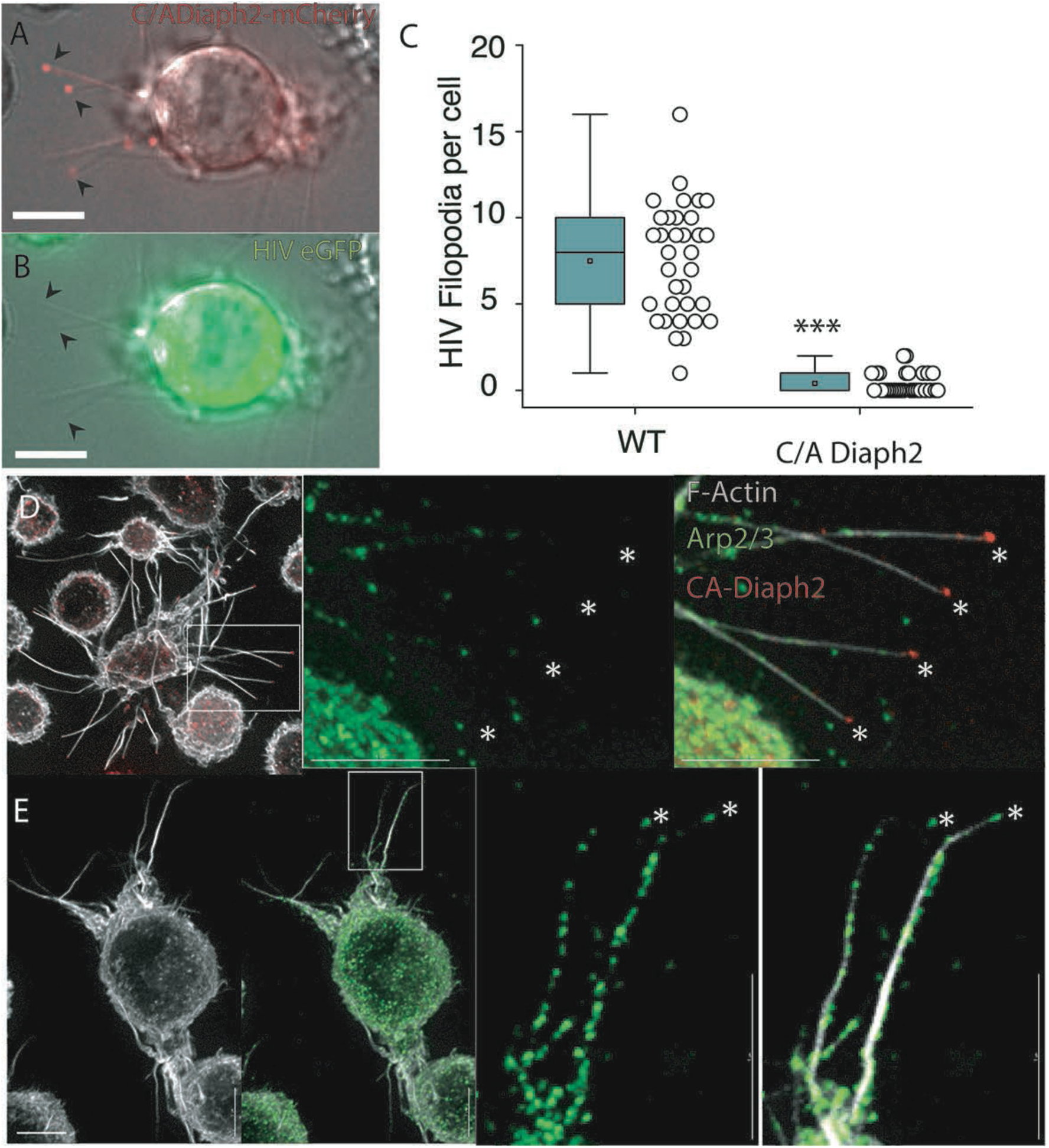
Constitutively active Diaph2 driven Filopodia are not associated with HIV buds. A & B. Live still images from supplement Movie S3; A. ^C/A^Diaph2-mCherry (red) positive cells, infected with B. HIViGFP (green). Note Diaph2 at the tips of filopodia are negative for HIV. Scale bars are at 5μm. C. Quantification of HIV positive filopodia per cell in HIV infected live cell cultures. HIV filopodia counts are derived from three independent HIV infections ***p<0.0001. D&E. Immunofluorescent Arp2/3 staining of D. ^C/A^Diaph2-mCherry (red) positive cells and E. Untreated cells. Asterisks highlight the terminal ends of filopodia that are either Diaph2 positive & Arp2/3 negative (for ^C/A^Diaph2-mCherry) or Arp2/3 positive (for untreated cells).

### HIV Gag can influence Arp2/3 dependent F-Actin pathways

Since HIV assembly at the PM is primarily driven by HIV-Gag, we turned to strategic Gag mutagenesis in an attempt to resolve the link of HIV assembly with cortical F-Actin structures. The HIV Gag mutant panel covered several well characterised mutants that could maintain HIV particle assembly and also binding to membrane Phosphatidylinositol (4,5)-bisphosphate (PIP2). Deletion of HIV Gag p6 and mutagenesis of the PTAP motif in p6, was used to block the recruitment of TSG101 and related ESCRT proteins involved in viral particle abscission (Fig. 4A-B). We also deleted the Nucleocapsid (NC) domain, as this has been previously proposed to mediate the interaction between HIV-Gag and F-actin [26]. However, since NC is required to facilitate higher order oligomerisation of Gag [27], we replaced NC with the Leucine Zipper (LZ) domain from the *Saccharomyces cerevisiae* GCN4 protein (Fig. 4C), as this rescues Gag oligomerisation and ensures particle assembly proceeds in the absence of NC [28]. Finally, given the enrichment of HIV buds on F-Actin structures with positive curvature, we further generated two HIV Gag Capsid mutants P99A and EE75,76AA (Fig. 4D & E), both of which inhibit Gag curvature at the PM but not high order Gag oligomerisation [29,30]. As Diaph2 cannot recruit HIV to F-Actin and depletion of Diaph2 actually enriched HIV-positive filopodia (Fig. S1), we utilised Diaph2^-ve^ cells and simply scored the number of filopodia per cell that were HIV positive for each viral Gag mutant. Using this approach, we observed no significant difference in viral filopodia when deleting p6, the PTAP motif in p6 or NC (Fig. 4F). However when using the P99A and EE75,76AA HIV capsid mutants (HIV curvature mutants), we observed Diaph2-depleted cells to not only lack any evident HIV buds at the PM but also their characteristic short filopodia. Instead, these cells resembled the lamellipodial phenotype observed in Diaph2^-ve^Arp2/3^-ve^ co-depleted cells (Fig. 4G-I). This is consistent with HIV Gag curvature mutants acting as dominant negatives for the Arp2/3-dependent short filopodial pathway. To further test that HIV curvature mutants were specifically interrupting Arp2/3 F-actin pathways and not broadly influencing all pathways that may lead to filopodial formation (eg. Formin-induced filopodia), we infected Diaph2^C/A^ cells with these mutants. In this setting we observed an ability of Diaph2^C/A^ to rescue filopodia formation (Fig. 4F; Movie S4). Thus filopodial pools nucleated by Arp2/3 are most affected by HIV curvature mutants and this further supports the hypothesis that HIV assembly primarily influences elements of Arp2/3 F-Actin nucleation pathway.

**Figure 4.**
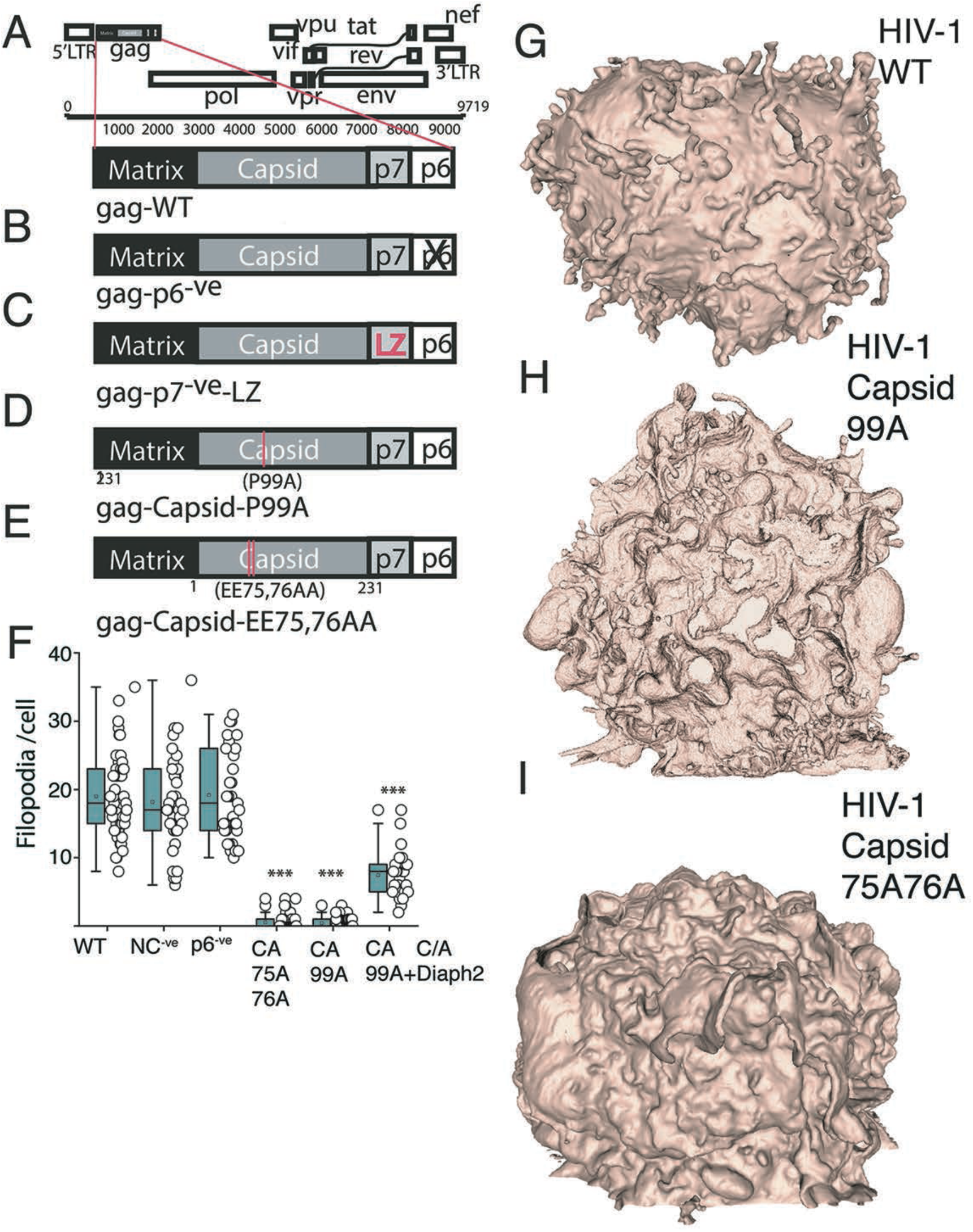
HIV Gag curvature mutants can impact Arp2/3 dependent cortical F-Actin. A-E HIV Gag mutants used. A. Wild type HIV Gag. B. HIV Gag late domain mutant with p6 deleted. C. NC deletion mutant with the Leucine Zipper (LZ) derived from *Saccharomyces cerevisiae* GCN4 to rescue Gag oligomerisation. D&E. HIV Gag curvature mutants D. P99A and E. EE75,76AA. F. Enumeration of filopodia per cell in Diaph2^-ve^ cell clones. ***=p<0.0001. G-H Representative FIB-SEM 3D rendered images of G. WT versus curvature mutants H. P99A and I. EE75,76AA. Data is derived from three independent HIV infections using each mutant.

### The HIV proteome reveals a GTPase node associated with Arp2/3 F-Actin regulation

HIV has been previously observed to incorporate F-Actin, various actin nucleators and numerous upstream/downstream regulators within virions [31-33]. Thus, we turned to mass spectrometry analysis of purified virions to observe the footprint of cytoskeletal proteins that are present at HIV assembly sites. For this analysis we also leveraged the three distinct F-Actin cell types generated above (i.e. long-filopodia, extensive short filopodia and large lamellipodia) as across each cell type they shared the feature of HIV buds being enriched in positively curved F-Actin structures. Using this approach we identified several Arp2/3 complex subunits, alongside two major Arp2/3 regulators, the Rho-GTPases Rac1 and Cdc42, as well as their interaction partner IQGAP1. These regulators were observed across all viral proteomes, irrespective of producer cell type (Fig.5 A&B). In addition to this F-Actin signalling node, HIV virions also acquired members of the integrin and cadherin families (Fig. 5 A-B; see nodes 3 and 4, respectively). These proteins, which are involved in cell-cell adhesion, are connected to the cortical F-actin cytoskeleton both physically and via signalling pathways [34]. Of interest was a depletion of the cadherin node in (Fig. 5 A; node 4), as well as an increase in Arp2/3 and Cdc42 content (Fig. 5 A; node 1) in virions produced by Diaph2-deficient cells. The latter observation is not only consistent with HIV assembly preferentially proceeding alongside short Arp2/3-dependent filopodia (as observed by FIB-SEM), but also suggests that these structures are dependent on Cdc42, which is a well known filopodial regulator [2]. Of note, the observed decrease of Arp2/3 components in virions produced in Diaph2 and Arp2/3 co-depleted cells (Fig. 5B, node 1) is both expected and consistent with depletion of these proteins at the cellular level (Fig. S3).

**Figure 5.**
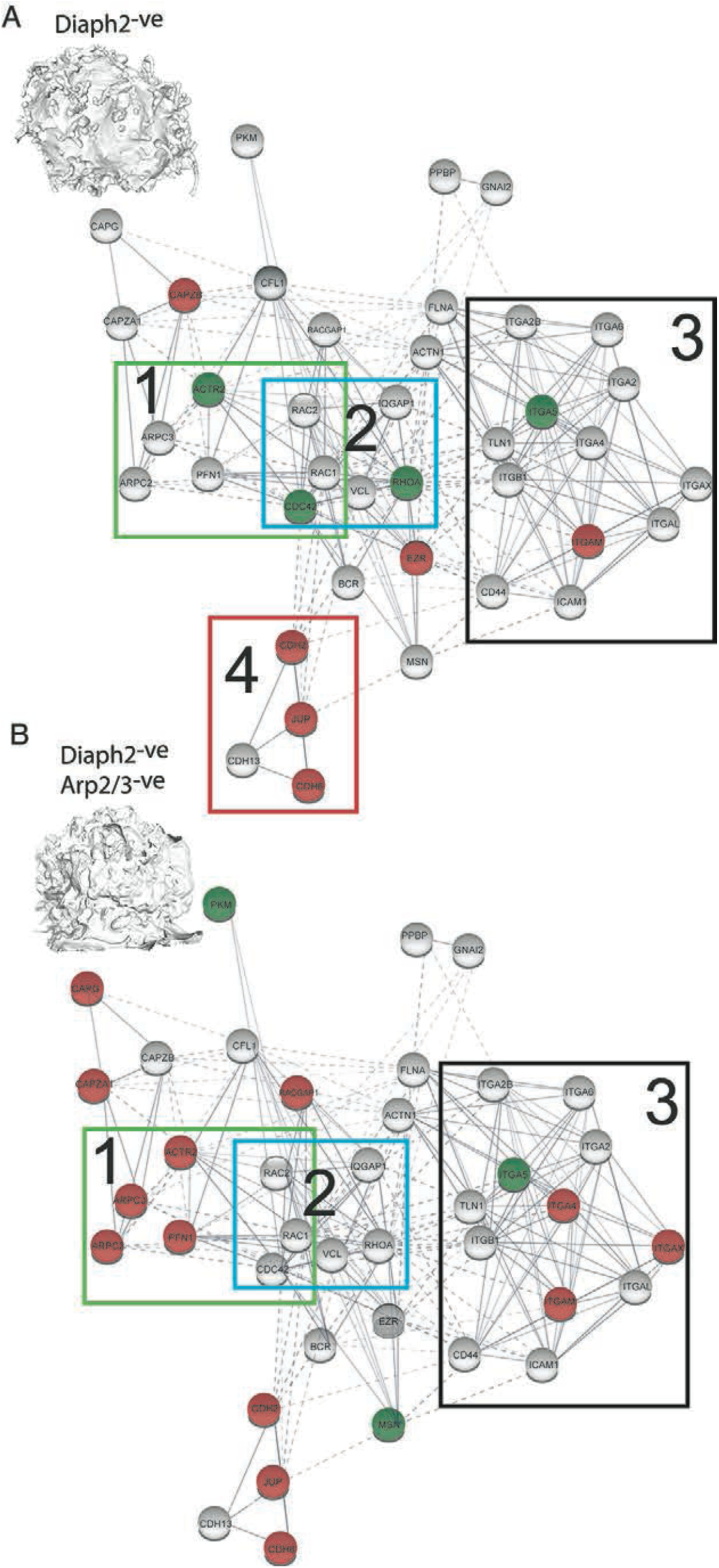
Cortical F-Actin regulators enriched at the final stages of HIV egress are revealed through HIV proteomics. A-B Viral proteome analysis of proteins associated with cortical F-Actin regualtion. Proteins with increased abundance relative to untreated cells are shown in green, whereas those with relative decreased abundance are shown in red. A. Virions produced in Diaph2^-ve^ cells. 1. Highlights the Arp2/3 complex node where the amounts of ACTR2 and Cdc42 are increased in virion proteomes. 2. Indicates a GTPase node in association with IQGAP1. 3. Highlights a node of integrin and related proteins. 4. Highlights a node involved in cadherin adhesion that is downregulated upon Diaph2 depletion. B. Virions produced in Diaph2^-ve^Arp2/3^-ve^ cells. Note that in 1. Arp2/3 components are predictably depleted compared to untreated cells, whilst in 2. the GTPase node and IQGAP1 remain unchanged.

### HIV exploits the Cdc42-Arp2/3 filopodial pathway to position virus at the leading edge of cell-cell contacts

Since Cdc42 is an important regulator of Arp2/3, a master regulator of filopodia, and it was incorporated at higher levels in virions from our Diaph2^-ve^ cells (more abundant short filopodia), we next targeted this protein for depletion. As a functional control, we targeted the homologous Rho-GTPase Rac1, best known for its role in lamellipodial regulation. We also investigated the scaffolding protein IQGAP1, which; i) is a binding partner and effector of both Cdc42 and Rac1 [35], and ii) plays an increasingly recognized role in actin cytoskeleton regulation [36], and iii) was consistently incorporated in virions in our experiments (Fig.5). While we succeeded in generating a viable Cdc42 homozygous knockout cell line using CRISPR/Cas9 (Fig. S4). Attempts at knocking out Rac1 led to multinucleated cell populations with reduced viability, which is consistent with previous reports of Rac1 being an essential gene [37]. To circumvent this, we partially depleted Rac1 by shRNA, and also generated a Wave2^k/o^ cell line (Fig. S4), since Wave2 is the main downstream effector of Rac1 in F-actin regulation [38]. While obtaining homozygous IQGAP1 knockout clones via CRISPR-Cas9 proved challenging, we were able to establish a line stringently depleted of IQGAP1 using shRNA (>99% depletion at the protein level, Fig. S3).

Initial Rac1 depletion via shRNA, revealed a greater frequency of filopodia in infected cells and secondly the generation of significantly longer and thicker filopodia when cells were infected (Fig. 6B vs A). Similarly, WAVE2^k/o^ cells infected with HIV had greater propensity to form filopodia (two-fold), and these were significantly longer and thicker compared to WT cells (Fig. 6C vs A and Fig. S2), but also uninfected WAVE2^k/o^ cells (Fig. S2). Together these observations suggest that HIV infection stimulates a pathway of filopodial formation that is unchecked in Rac1^-ve^ and WAVE2^k/o^ cells, where the lamellipodial F-actin arm is disabled. Given the known role of Cdc42 in filopodia formation and its competing nature with the Rac1 pathway, we turned our attention to this Rho-GTPase. Importantly, Cdc42^k/o^ cells were devoid of filopodia and coalesced cortical F-Actin into prominent lamellipodia, with no evident influence on F-actin when cells were HIV infected (Fig. 6D). Since IQGAP1 has been previously reported to articulate Cdc42 signaling to the cytoskeleton [36], we also assessed the role of this regulator in the filopodial context. IQGAP1-deficient cells displayed a collapse in filopodial lengths (Fig. 6E), with maintenance of HIV at the leading edge of remaining filopodia, similar to that observed in Diaph2-depleted cells. We therefore conclude that IQGAP1 can influence filopodial networks but, like Diaph2, is not required for the seeding of filopodia. To summarize our combined observations from mass spectrometry, gene silencing and high-resolution imaging, reveal that HIV infection augments a pathway of filopodia formation, and this is most evident when the lamellipodial regulators are inactivated. In contrast, removing Cdc42 completely blocked filopodia formation in a manner similar to Arp2/3 and Diaph2 co-depletion, whereas depletion of IQGAP1 or Diaph2 led to shorter filopodia. Together, our data indicates that HIV-assembly hijacks a cellular pathway that is dependent on Cdc42-Arp2/3 F-actin nucleation for filopodial seeding and IQGAP1/Diaph2 for filopodial elongation, in order to position itself at the tips of long filopodia.

**Figure 6.**
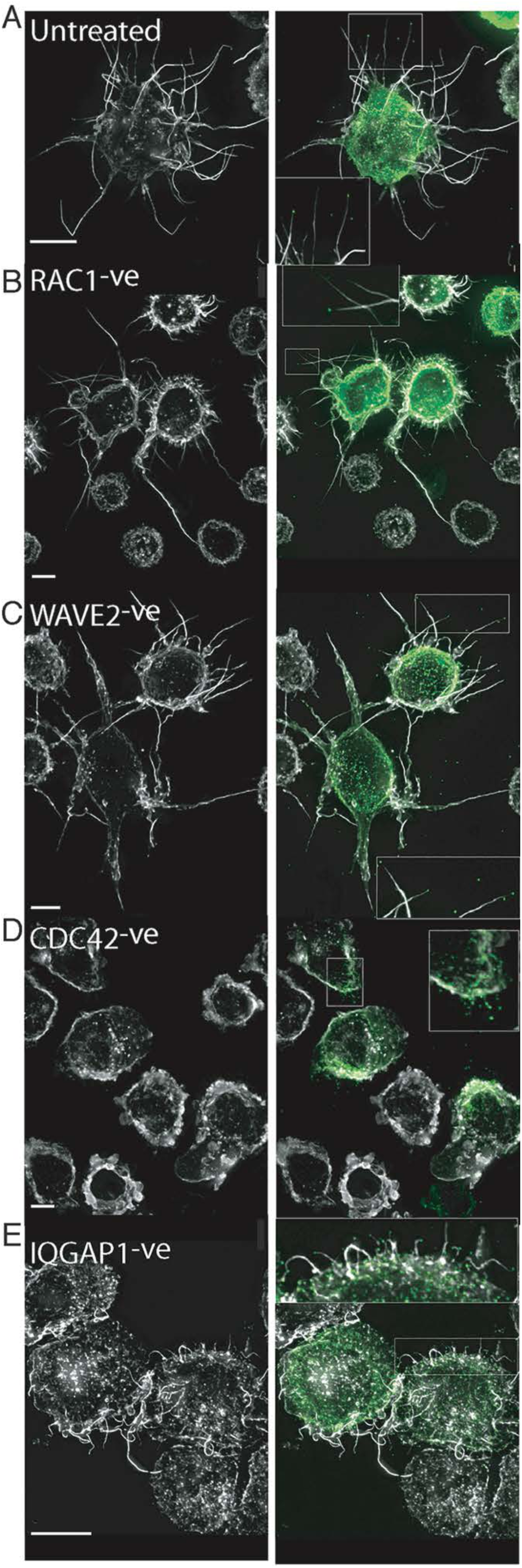
HIV infection and its influence on cortical F-Actin. A. From viral proteomics, we identified a common node of actin regulators sassociated with Arp2/3-dependent filopodia and lamellipodia. Through shRNA depletion or CRISPR-Cas9 knockout, we generated clonal cell populations depleted of various actin regulators. B-C Lamellipodial regulators. B. Rac1^-ve^ and C. WAVE2^k/o^. D. Cdc42^k/o^ (filopodial regulator). E. IQGAP1^-ve^. A. Represents the untreated control. All cells were infected with HIV iGFP (green) and then counterstained with phalloidin Alexa-647 (white). All scale bars are at 5μm. Inset magnifications reveal HIV at the leading edge of filopodial structures. Note in Rac1^-ve^ and WAVE2^k/o^ images the extensive filopodial networks only present in HIV infected (green) cells (see Fig. S2 for formal quantification).

### HIV cell-cell transfer is dependent on an intact Cdc42-IQGAP1-Arp2/3 pathway

Given the continuum of phenotypes observed in our abovementioned observations, we tested their impact on the late stages of the viral life cycle in the context of viral spread. For free virus release, we enumerated HIV particles accumulating in the supernatant as a measure of budding. As HIV spread can also proceed through direct cell-cell contacts, we further tested the ability of HIV to spread cell to cell by coincubating infected donor cells with permissive target cells. Using these approaches, we could determine if the generic lack of a cortical F-Actin structure or a specific F-Actin pathway is essential for HIV budding and/or cell-cell transfer.

Disruption of either the Rac1-WAVE2 pathway (lamellipodia) or Cdc42-IQGAP1 pathway (filopodia) both impaired free HIV budding, as indicated by significantly lower viral particle counts in the supernatant from cells depleted of these regulators, compared to untreated cells (Fig. 7 B). Furthermore, lack of budding was not associated with decreased viability of each cellular clone or lack of HIV Gag expression (Fig. S3 H & I respectively). However, impaired release of free HIV did not predict outcomes for cell-cell HIV transfer. For cells with disabled Rac1/WAVE2 (Rac1^-ve^, Wave2^k/o^ cells) cell-cell HIV transfer persisted (Fig. 7 A&B), despite the decreased free virus budding ability. In contrast, disruption of the Cdc42/IQGAP1 axis (Cdc42k/o and IQGAP1-ve cells) impacted both HIV budding and cell-cell transfer (Fig. 7 A&B). These observations suggest that while normal actin dynamics are important for free virus release, Cdc42 and IQGAP1 are specifically required for cell-cell HIV transfer, whereas Rac1/Wave2 are not. To further test this hypothesis in the setting of primary CD4 T cell targets, we focussed cell-cell transfer assays with disruption of the Rac1-WAVE2 pathway versus disruption of Cdc42-IQGAP1. In this setting, we further tested the efficiency of cell-cell spread by limiting dilution of the infected donors into primary CD4 T cell co-cultures. Using this approach, we observed almost complete loss of cell-cell HIV transfer in Cdc42^k/o^ and IQGAP1^-ve^ clones, whereas cell-cell transfer persisted in Rac1^-ve^, Wave2^k/o^ clones, albeit slightly lower than in WT cells (Fig. 7C). In cells lacking filopodia (CDC42 and IQGAP1), one immediate mechanism for lack of viral transfer could be the culmination of a limited contact capacity with the cells immediate microenvironment. To test this hypothesis, we enumerated accumulative cell to cell contacts (Fig. 7D) and later target cell engagement (Fig. 7E) in wild type, IQGAP1^-ve^ and WAVE2^k/o^ cells. IQGAP1^-ve^ cells were observed to have significantly lower overall contacts and also engaged fewer cell targets. Whilst this addresses the lowered ability of cells without filopodia to participate in cell-cell transfer, it does not address the address the paradox of persistent cell-cell HIV transfer, despite reduced viral budding in cells with augmented filopodia. To resolved this further, we observed later interactions WAVE2^k/o^ cells where filopodial networks are augmented following HIV infection. Filopodia initially persisted in early cell-cell conjugates, yet we routinely observed collapse of filopodial networks immediately preceding VS formation and HIV-GFP transfer to the opposing target cell (Fig. 7 F and Movie S6). As a surrogate of filopodial activity, we quantified this as membrane complexity through calculation of cellular circularity. In this setting, cells with extensive filopodial networks were observed to have low circularity, whilst cells with no filopodial activity were observed to have high score in circularity. Measurements of Gag-polarisation over time then established measurements of the seeding of the VS and release of GFP into the neighbouring target was used to mark the final stage of VS maturation that culminated in viral transfer. Using this quantification in the representative live cell movie acquisitions (7 F &G), we observe cells engaged in cell-cell contact to approach a circularity of 1 (i.e. Cells collapsing their filopodial networks) just prior to the final stages of viral transfer, as marked intitially by Gag polarisation and then subsequently observed in cytoplasmic transfer of GFP to the neighbouring cell (7G).

**Figure 7.**
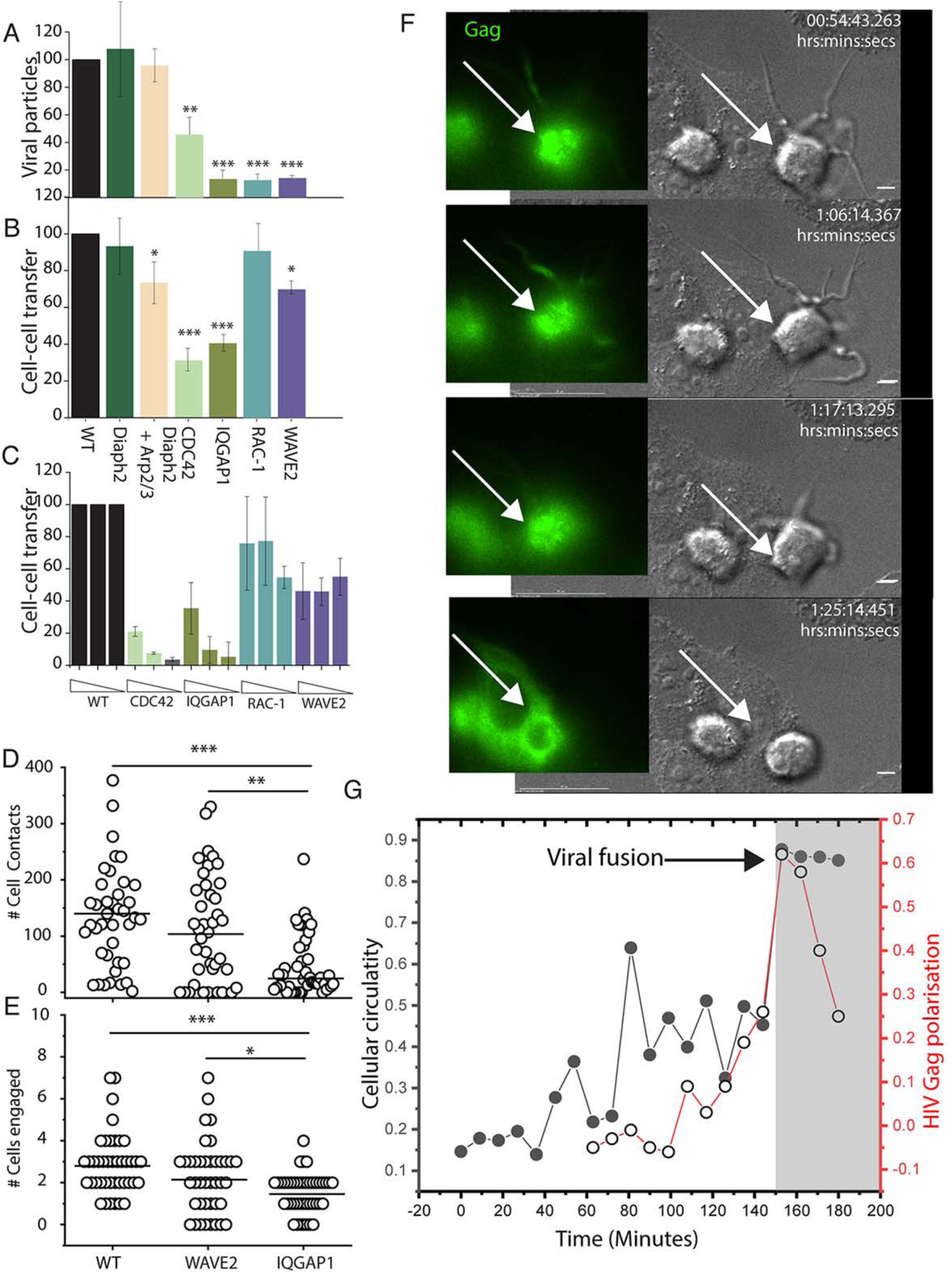
HIV spread is dependent on Cdc42 and IQGAP1. A. All cells are infected with pHIVNL43iGFP and normalised to 5% infection on day 3. After normalisation, cell supernatants are collected over a 24 hour period and GFP positive HIV particles are spinoculated onto 96-well glass plates coated with poly-L-lysine. Absolute viral particle counts are determined by high-resolution fluoresecence microscopy per 4 fields of view. Herein the data is presented as a relative count [(virion count/virion count in WT control)*100]. B. HIV infected cells as normalised in A. are then co-cultured at a ratio of 1:5 with HIV-permissive TZMBl targets. C. Infected cells are co-cultured with primary CD4 target T-cells at limiting dilutions. Dilution steps correspond to 5%, 1% and 0.2% infected cells in the donor population. Exposure to virus from infected cells is limited to 24 hours, after which an entry inhibitor BMS806 is added to prevent further viral spread. A-C. Data indicates the mean and standard deviation from 3 independent experiments. In C, primary recipient CD4 T cells were sourced from independent blood donors. D) The cumulative number of contacts between each infected donor cell and any uninfected target cells (TZM-bl) over 3 hours. E) same as D but only the first contact with each distinct target cell is counted. F) Representative example of a time-lapse series from D-E. Cells are infected with HIViGFP, allowing real-time visualization of Gag. G) At the virological synapse, donor-cell circularity was used to enumerate lack of membrane protrusions (ie. Lack of filopodia) in parallel with Gag polarization. In grey shading are the time points where GFP cytoplasmic transfer (ie. Viral fusion) is observed. * p<0.01, **p<0.001, ***p<0.0001.

**Figure 8.** Proposed model of spatio-temporal regulation of F-Actin during HIV egress. A. With probing and tethering activity corrupted by HIV buds, the F-actin structures at this phase enable pre-synpatic events (cell contacts and initial adhesion). B. Following cell-cell engagement, the viral synapse matures. This leads to two important outcomes. Firstly, similar to that observed at the immunological synapse [72] Cdc42-F-Actin activity is altered and transitions from filopodial biogenesis to cell-cell adhesion required for HIV release (as observed when cells collapse filopodia just prior to HIV fusion). During this change we hypothesise IQGAP1 being directly involved through providing a scaffolding center needed temporally coordinate binding not only to HIV Gag, but also providing feedback to other binding partners including Cdc42 and TSG101. Whilst the observations herein and recently by others [41], initially supports this model, future work will be key in understanding the switching nature of IQGAP1 and how F-Actin and CDC42 influences its role in the final stages of viral absciscion and transfer.

## Discussion

The corruption of cortical F-Actin by HIV has remained elusive, with evidence both for and against its role in budding and cell-cell viral spread. Through systematic depletion of various F-Actin regulators, combined with viral mutagenesis and high-resolution imaging, we were able to illuminate the intersection of HIV egress with cortical F-Actin and conclude this is primarily associated with the Cdc42-IQGAP1-Arp2/3 pathway.

Our primary aim herein was to understand how HIV egress was spatiotemporally connected to a continuum of cortical F-actin structures that dynamically regulated in leukocytes. Whilst many prior studies have mapped F-Actin pathways in cell-free systems, the challenge herein was to map F-Actin pathways and how they influenced not only the live virus, but also in a cellular & cytoskeletal setting that was consistent with that of the immune system. Whilst many F-Actin regulators are common across cells, F-Actin regulation in leukocytes is unique and enables a rapidly changing canvas of F-Actin polymers to coordinate their roles in the immune response. Processes such as chemotaxis, promisicous cell-cell scanning and later stable cell-cell contacts are all processes dependent on dynamic cortical F-Actin. So in discussion of the how F-Actin influences HIV egress we need needs to to frame the discussion into two distinct events. Firstly, HIV’s time at the leading edge of protruding F-Actin structures (target selection) and secondly, when infected cells engage in longer stable contacts (target engagement).

At the leading edge of F-actin, we observed HIV buds to be enriched where cortical F-actin structures induce strong positive curvature. However, curvature and/or formin activity alone were not sufficient to position HIV at the tips of filopodia. Thus, other actin regulators associated with membrane curvature must be involved. Our observations herein that HIV infection specifically enhances Cdc42-Arp2/3-dependent filopodia, supports a mechanism of action that corrupts this pathway of actin nucleation. While curvature provided by HIV during budding could itself drive filopodia formation (e.g by direct recruitment/activation of Cdc42/Arp2/3 [39,40], our observations support hijacking of a pre-existing pathway dependent on curvature. We base this reasoning three-fold. Firstly on uninfected myeloid cells there are similar long filopodia (albeit uncapped with HIV). Secondly most filopodia in infected cells are HIV-capped [9]. If HIV would provide an independent mechanism of filopodia formation, both structures would be expected to coexist. Thirdly and finally, HIV curvature mutants had a dominant negative effect on all filopodia, indicating i) a functional overlap of the viral and cellular pathways of filopodia formation, and ii) a critical role of Gag in hijacking of these structures. Recently, Sabo and colleagues observed HIV Gag to directly interact with IQGAP1 [41]. This observation is consistent with our observations herein at several levels. Firstly, as IQGAP1 binds to Cdc42 and stabilizes it in its active conformation to drive filopodia formation [36]. Secondly, as IQGAP1 also facilitates assembly of multiprotein complexes that spatially link Cdc42, Arp2/3 and formins [42]. This is consistent with how HIV is associated with a filopodial structure that is firstly nucleated by CDC42, but secondly elongated by the formin Diaph2. Finally, given the role of positive curvature in filopodia biogenesis, consumption of IQGAP1 by HIV Gag into an area of neutral curvature, is consistent with HIV Gag curvature mutants acting as dominant negative constructs for CDC42-Arp2/3 filopodia. Whilst the resolution of pathways that gives birth to HIV filopodia are now becoming clearer, the role of this hybrid viral and cellular structure now needs discussion. Typically, viral-cellular membrane events are associated with viral release. In contrast, we observed HIV’s association with F-Actin to be inhibitory to HIV budding. The most evident is when HIV filopodial networks are formed during Rac1 and Wave2 depletion. In that setting, the removal of Rac1 would not only bias signalling to CDC42, but with IQGAP1 recruited by HIV Gag, CDC42 would be maintained in an active GTP bound state [43]. This is entirely consistent with its role in filopodial formation but also its augmentation by HIV Gag-IQGAP1 during infection. Whilst arrest of viral HIV buds in a F-Actin structure may be a function of distance that this structure projects immature HIV from the membrane, the role of the virus in this setting is seems counter intuitive at two levels. Firstly, a virus that cannot undergo absciscion at the membrane cannot subsequently mature [44] and as such this cannot contribute to viral spread. Secondly, it is well known HIV particles bud from infected cells. To reconcile both observations, we hypothesise leading edge structures positive with HIV can indeed indirectly contribute to cell-cell spread and that only a proportion of HIV buds at the membrane engage in this process. So in this setting, the virus at the leading edge of protruding F-Actin structures may indeed be hard-wired to coordinate the initial pre-synaptic contacts. Yet as cell-cell contacts mature, HIV’s relationship with F-Actin we support observed to be reconfigured, as the inhibitory mechanisms observed during pre-synaptic events are overcome during cell-cell viral transfer. So to conclude, HIV’s time at the leading edge of a cell we support to function primarily in pre-synaptic events. In this context, HIV’s corruption of the leading edge does not contribute to viral release but directs initial cell-cell contacts.

Shadowing and highjacking the Cdc42-IQGAP1-Arp2/3 actin regulatory axis is not a unique feature of HIV. Nascent viral buds have also been observed at the tips of filopodia-like structures for other types of viruses [45-50], and numerous intracellular pathogens are known to exploit Rho-GTPases and the unique ability of the Arp2/3 complex to promote formation of specialized cortical F-actin membrane protrusions that facilitate cell-cell infection spread [51-55]. Similarly, IQGAP1 is a prominent target of microbial manipulation and this is closely related to its ability to modulate the actin cytoskeleton (reviewed in [56]). Several viruses bind IQGAP1 either directly (via interactions with the viral matrix protein) or indirectly (via common binding partners) and this has important consequences for viral assembly, budding and/or pathogenesis [11,47,57-60]. For HIV, recent studies by Sabo and colleagues [41], have observed biochemically IQGAP1 interactions with NC & p6 elements of HIV Gag. In this study they support a role for IQGAP1 in negative regulation of HIV Gag trafficking and subsequent HIV-budding [41]. Whilst our observations readily support a role for IQGAP1 in influencing HIV budding, we did not observe this to be a consequence of negative regulation of Gag trafficking and docking to the membrane. For instance near complete removal of IQGAP1 did not increase viral budding and egress, but rather led to inhibition thereof. In light of our observations herein and those recently published [41], we would conclude that as IQGAP1 is a scaffolding protein with many binding partners, it is not surprising that HIV Gag has many different fates depending on each cell type it is expressed in and what functions that cell type maybe engaged over the time the cells were sampled. Importantly, we do readily support a role for IQGAP1 in the viral life cycle and this readily supports recent observations by this team.

In terms of HIV spread, free viral particle release was susceptible to inhibition of both lamellipodial and filopodial regulators. In contrast, cell-cell spread of HIV was mainly dependent on Cdc42 and IQGAP1 but showed tolerance to depletion of Rac1/Wave2, despite a similar impact of all regulators on free virus budding. Together, these observations suggest that long filopodia specifically contribute to cell-cell HIV spread, whereas lamellipodia are less important for this process. This is consistent with findings from other enveloped viruses where filopodia have been associated with the ability to mediate cell-cell viral transfer [61-65]. Indeed, we have previously observed HIV-Filopodia to mediate hundreds of contacts per hour between relevant primary HIV target cells, with filopodial activity often preceding VS formation [9]. The latter is in agreement with previous observations that filopodia and/or dendrites may commonly serve as precursors for biological synapses [66-70]. It has also been proposed that, as synapses mature, filopodia must be “supressed to allow a smooth and broad cell-cell interface” [70]. This is consistent with our observations herein that early HIV donor-target cell contacts are characterized by abundant filopodial activity, whereas filopodia are often lost during late stages of VS formation, when polarization and transfer of Gag is most evident. These remarkable cell-shape changes reveal that VS progression involves extensive cytoskeletal remodelling and suggest a clear switch of actin-manipulation strategy as the synpase matures. Loss of filopodia likely requires inactivation of Cdc42, and this could be important to allow synchronised budding and large-scale viral release at the VS, in a similar manner to how Cdc42 inactivation promotes mechanistically analogous (i.e. ESCRT-dependent) abscission events during late stage cytokinesis. Thus, IQGAP1 may serve as a scaffolding center that enables coordination and cross-signaling of F-actin remodelling and abscission events during both cytokinesis and the VS. However, as with other synapses, maturation of the VS is a temporally and mechanistically complex process, and further studies will be required to fully elucidate the mechanisms involved.

Overall, we propose that HIV has evolved to highjack a specific node of F-actin regulation that positions outgoing virus at the leading edge of cortical F-actin structures. Since filopodia play an important role in scanning of the microenvironment and mediating immune cell interactions, their corruption is beneficial to the virus because it biases the first line of cell-cell contacts towards HIV spread, e.g. by providing enhanced adhesion and specificity to CD4+ target cells. However, as these contacts form and the VS matures, the relationship of HIV with F-Actin changes and observations herein support this change to facilitate final viral release. We conclude that manipulation of the Cdc42-IQGAP1-Arp2/3 actin regulatory node is essential for corruption of the cells leading edge to facilitate cell-cell HIV spread, and observations support a key role for IQGAP1 also during later cell-cell contact. Moving forward, greater spatiotemporal resolution of this latter events is needed and will give further insight into why many pathogens like HIV have evolved to interact with IQGAP1 and its binding partners.

## Materials and Methods

### HIV Plasmid constructs

Plasmid constructs used herein are all based on CCR5 using pNL43AD8^ENV^, unless otherwise indicated (Table SI). All details regarding GFP carrying HIViGFP and HIViGFP^ENV-ve^, including the insertion of GFP in Gag polyprotein, have been previously described [9]. HIV nucleocapsid mutant (gag-p7^-ve^-LZ) was generated using a two-step cloning strategy wherein a 861 bp fragment carrying the HIV Capsid domain, P2 spacer region and the leucine zipper domain of yeast transcription factor GCN4 (herein LZ) was amplified from plasmid pRR546, a kind gift from Dr A Rein (NCI, USA)[28]and shuttled into HIViGFP using *XbaI/ApaI* cut sites. The intermediate plasmid (HIV^LZ1stGen^) thus generated had most of the NC domain replaced with LZ domain except for a 75 bp region proximal to the p6 domain. To remove this fragment and have the LZ domain contiguous with the p6 domain, primers were designed to amplify a 760 bp fragment from HIViGFP containing the p6 domain and Pol region proximal to Sbf1 restriction site while simultaneously introducing an *Apa1* restriction site at the 5’end. The resulting amplicon was then shuttled into HIV^LZ1stGen^ using *ApaI/SbfI* cut sites to generate gag-p7^-ve^-LZ. Capsid mutant constructs gag-Capsid-EE75,76AA and gag-Capsid-P99A were generated by site directed mutagenesis wherein the reverse primers were designed to include the desired mutations. For gag-Capsid-EE75,76AA, a 320 bp fragment downstream of GFP was amplified from HIViGFP with the mutagenesis primers carrying the respective mutations (EE75,76AA) and re-cloned into the parental construct using *XbaI/SphI* unique sites while for gag-Capsid-P99A a 370 bp fragment downstream of GFP was generated with the mutagenesis primers carrying the point mutation, P99A, and shuttled back using *XbaI/SpeI* sites. HIV p6 deletion mutant gag-p6^-ve^ was synthesised by cloning a 1.5 kb fragment from plasmid L1-term, courtesy Dr Eric Freed, NIH, USA, into HIViGFP using *SpeI/SbfI* restriction sites. The construct L1-term carries a stop codon at the start of p6 domain of Gag in pNL43 and has been described in detail elsewhere {11805336}. pHIVNL43IRESeGFP (Courtesy of Dr Paul Cameron, Doherty Institute, Melbourne) expresses eGFP in the Nef open reading frame followed by an IRES element and the intact HIV Nef open reading frame. All of the above clones were sequence verified and transfected into HEK F293T cells (Invitrogen) as previously described[9].

### Virus production and infections

In order to examine Gag mutations in context of infectious virus, we generated viruses capable of single round of infection by ‘rescuing’ HIViGFP Gag mutants with a 2^nd^ generation lentiviral packaging construct psPAX2 (courtesy of Dr Didier Trono through NIH AIDS repository). The latter construct expresses wild type HIV Gag and Gag-Pol under a CMV based promoter and enables HIV genomes encoding the indicated Gag mutants to be packaged into viral particles (consisting of both mutant and WT Gag proteins) and importantly entering and infecting cells in a single round of infection. As psPAX2 only supplies WT Gag and Gag-Pol at the protein level, following infection, only the products of the mutant HIV genome are expressed (i.e. only the Gag mutant protein). To generate ‘rescued’ HIViGFP Gag mutants viral stocks for U937 infections, the mutant constructs were co-transfected with psPAX2 and also the VSVg plasmid pMD2.G (Addgene; courtesy of Dr Didier Trono) at a molar ratio of 2:1:1 to increase the infection rate. Transfections were done in HEK F293T cells using polyethylenimine (at 1mg/ml, pH 7.0) as described previously [9]. For cell to cell transfer assays, pHIVNL43IRESeGFP was used and was produced by polyethylenimine transfections of HEK F293T cells. TZM-bl indicator cell line (courtesy of the AIDS reagent repository) was used for virus titering and analysis as previously described[9].

### Lentiviral constructs and production

#### Diaph2^C/A^ mutant construct

Lentiviral plasmids expressing the constitutively active form of Diaph2 (pLVXdeltaDADmcherry) fused to mCherry was synthesised using the *XhoI/ApaI* restriction sites. The 30 bp autoregulatory domain (DAD) of Diaph2 with the consensus sequence DET(G/A)(V/A)MDXLLEXL(KIR/Q)X(G/A)(S/G/A)(A/P) spans aa 1051-1081 at the C-terminus and was removed from Diaph2 cDNA during cloning. Forward and reverse primers carrying the above restriction sites were designed to amplify a 3kb fragment from Diaph2 MGC human cDNA clone (Dharmacon) and the amplicon encompassing the entire cDNA sequence minus the DAD domain was cloned into pLVX-mCherry-N1 (Clontech) to generate lentiviral expression vector pLVXdeltaDADmcherry. The plasmids were sequence verified and lentiviral particles were then generated using the helper plasmid, psPAX2 and pMD2.G as previously described[9].

#### Lentiviral shRNA vectors

shRNA sequences for each gene target were obtained from The RNAi Consortium (TRC) library database and sequences with high adjusted scores (2 shRNA sequences per gene) were selected for synthesis (Table SII). Oligonucleotides (both sense and antisense) carrying *Age1/EcoR1* sticky ends, hairpin loop sequence and shRNA sequence were synthesised (IDT technologies) and annealed oligos cloned into pLKO.1 TRC cloning vector (Addgene #10878) using the unique *Age1/EcoR1* sites. A pool of two shRNA plasmids per gene was then packaged into lentiviral particles using psPAX2 and pMD2.G as previously described[9]. Alternatively and where indicated, shRNA plasmids (pool of three shRNA plasmids per target) were obtained directly from Santa Cruz.

#### Lentiviral CRISPR vectors

The SpCas9 and guide RNA (gRNA) CRISPR components were both expressed from the one-vector lentiviral system “lentiCRISPR_v2” {25075903} (Addgene #52961). gRNA sequences for target genes were designed by submitting the sequence of an early exon (common to all isoforms) into the CRISPOR prediction tool{Concordet, 2018 #1425}, so that frameshifts in this region would result in at least 70% loss of native protein sequence. gRNAs were selected to meet the tool’s specificity-score requirements and to have at least 4 base pair mismatches with any other exon in the human genome (Table SIII). gRNA oligos were ordered from IDT-Technologies and cloned into the lentiviral vector as outlined by Zhang and colleagues {25075903} (http://genome-engineering.org/gecko/wp-content/uploads/2013/12/lentiCRISPRv2-and-lentiGuide-oligo-cloning-protocol.pdf).

### Cell culture, genetic modification and infection

Monocytic cell line U937 (ATCC® CRL1593.2™) cultured in RPMI (Thermoscientific) supplemented with 10% FCS (Invitrogen) was used in all experiments, unless otherwise stated. The identity of the U937 cell line used throughout this study was verified by microsatellite profile analysis by a NATA accredited third party institution (Garvan Institute, Sydney) and was confirmed to match the U937 cell line in the ATCC and SMZ databases. The cells were maintained at a density of around 0.5 - 1 × 10^6^ per ml and passaged every 3 to 4 days. Although HEK F293T and TZMbl cells lines were not verified by microsatellite profile analysis, unique cellular resistance and phenotypic properties of cells were used to ensure purities were maintained. For HEK F239T, this was G418 resistance and the ability to produce HIV viral particles post transfection with the vectors used herein. For the TZMbl line, the ability to be infected by both CCR5 and CXCR4 tropic HIV strains in addition to their ability to produce luciferase and/or ß-galactosidase post infection was used to ensure purities were maintained. All cell lines used in this study were tested to be free of mycoplasma using Mycoalert (Lonza).

#### shRNA depletion

For shRNA knockdown in U937 cell lines, cells were transduced with lentivirus stocks expressing respective shRNA at an MOI of 1.0 and transduced cells selected by passaging in media containing puromycin (2µg/ml) (Invitrogen) or hygromycin (400µg/ml) (Invitrogen) for 2 weeks. For double knock down cells, transduced cells stably expressing single gene knockdowns were infected with lentivirus carrying the shRNA for the second target and transduced cells selected by addition of both puromycin and hygromycin to culture media at the above mentioned concentrations.

#### CRISPR gene knock outs

For gene-editing using CRISPR vectors, U937 cells were transduced at a MOI of 0.5, and selected in puromycin-containing medium (2µg/ml) for 7 days.

#### Clonal sorting, depletion and knock out confirmation

Clonal populations for the above-mentioned transduced cell lines were generated by single cell sorting using ARIA flow cytometer (BD Biosciences) into complete RPMI media that was pre-conditioned for 24 hours with the U937 cell line cultured at 2×10^5^ cells per ml. Approximately 30% of single cell colonies survived and were grown to confluency. Initially 5 clones were selected for live imaging and F-Actin phalloidin staining to reveal any readily observable changes in cortical F-Actin architecture (e.g. Loss of filopodia). Clones were then verified for shRNA depletion and gene knockdown at the protein level was verified by western blotting of cell lysates using the following antibodies: anti-Arp2 rabbit, polyclonal (Santa Cruz; SC-15389; Lot L-2109), anti-Arp3 mouse, monoclonal (Abcam; AB49671; Lot GR450638), anti-Diaph1 rabbit, polyclonal (Bethyl laboratories; A300-077A; Lot A300-077A-2), anti-Diaph2 rabbit, polyclonal (Bethyl laboratories; A300-079A; Lot A300-079A-2), anti-Diaph3 rabbit, polyclonal (Protein tech; 14342-1-AP; Lot 5399), Anti-Rac1 monoclonal (Protein Tech; 66122-I-Ig; Lot 10011346) and anti-IQGAP1 monoclonal (SantaCruz; sc-376021; Lot H1417) antibodies. In the absence of working antibodies to verify protein target depletion, RT-Q-PCR was performed on FMNL-1 (HS00979762_m1) using Applied Biosystems TaqMan Gene Expression assays and fold change depletion in the shRNA depleted cells was determined by comparison to scrambled shRNA transduced cell controls. In supplementary Figure S3, raw data (immunoblotting of cellular lysates) is presented on the clonal cells primarily used in this study. Where possible, CRISPR/Cas9 was used to confirm shRNA depletion phenotypes and also to generate cells where the target protein expression was knocked out. Successful gene editing in clonally expanded populations was confirmed by genomic PCR and DNA sequencing with a pair of target-specific “surveyor” primers designed to amplify the gene region targeted by the gRNA (all surveyor primers used are listed in Table SIII). Genomic DNA extraction from CRISPR-edited cells was performed as described in {Guschin, 2010 #129}. Briefly, 1 × 10^6 cells were pelleted, the cell pellet resuspended in 100μL QuickExtract solution (Lucigen #QE09050), transferred to PCR-tubes and incubated in a thermocycler for 15 minutes at 68°C, followed by 8 mins at 95°C and then held at 4°C until further use. Of this cell-lysate solution, 2μL were used as template for genomic PCRs in a 50 μL reaction with Velocity DNA Polymerase (Bioline#21098) following the manufacturer’s instructions. DNA Sanger sequencing was performed by a NATA-accredited core facility (Garvan Institute, Sydney). Sequencing data was analysed using Sequence Scanner Software v2.0. Note that in the context of this work, the term knockout (k/o) refers to cellular clones where both gene copies have been edited in a way that results in loss of native protein expression due to introduction of frameshifts or premature stop codons in the protein coding sequence. For representative sequencing data of Cdc42^k/o^ and Wave2 ^k/o^ cell clones, see Fig. S4.

### Quantitative cell to cell virus transfer assays

Primarily viral transfer assays utilised various U937 clones with shRNA depletion or gene knock outs where indicated. Donor cells were infected at an MOI of 1.0 with VSVg pseudotyped pHIVNL43IRESeGFP by spinoculating at 1200 x g for 1 hour at 18°C. 48 hours post infection, the proportion of infected cells was enumerated by measuring GFP expression via flow cytometry.

Unless otherwise stated, viral transfer assays used the HIV permissive HeLa cell line TZM-bl cells. 5000 infected donor cells normalised to 40% infection were added to 20,000 TZM-bl cells (labelled with “Thermofisher-C34565: CellTracker Deep Red dye” and seeded on the previous day) in a flat bottom 96-well plate (donor to target ratio = 0.25, final well volume = 200 µl). Viral transfer was stopped after 18 hours coculture by adding the gp120/ CD4 inhibitor BMS-378806 (Selekchem #S2632) to a final concentration of 10 µM. At 48 hours coculture, donor cells were removed by washing twice in PBS. In all cases, the number of infected recipient cells (GFP and Alexa647 double positive, single cells) were enumerated by fluorescence microscopy using the high content CYTELL imaging platform (GE Healthcare). An average of 25,000 cells were analysed per condition by acquiring multiple fields of view.

In virus transfer assays that used primary CD4+ T cells as targets, primary CD4+ T cells were isolated from whole blood. Healthy donors were consented under St Vincent’s Hospital ethics #HREC/13/SVH/145. Briefly, blood from each consenting donor was collected into approximately 10x 9ml ACD-B Vacuette tubes (Greiner, Cat# 455094; Lot#A180437V). Blood was then pooled and diluted I in 2 with sterile PBS and then subsequent CD4 T cell isolation proceeded as previously described[9]. Following isolation, cells were activated using T Cell TransAct™ (Miltenyi Biotech) for 24 hours according to manufacturer’s instructions. CD4+ T cell activation was measured by upregulation of CD69 (Becton Dickinson Clone L78; Lot 41209) and CD25 (Becton Dickinson Clone 2A3; Lot 25912) surface marker expression using flow cytometry. 10^5^ primary activated CD4+ T cells were added per well to a U-bottom 96-well plate in 100µl of RPMI supplemented with 10% fetal calf serum. Following extensive washing and normalisation of infection to 5%, 5000 donor cells per 50µl of media were serially diluted at steps of 1in 5 dilution and added to recipient cells in a final co-culture volume of 150µl. Four days post co-culture, cells were thoroughly resuspended and added to wells of a 96-well flat bottom tissue culture plate that had been previously coated with poly-L-Lysine as per manufacturer’s instructions (Sigma-Aldrich). As mentioned above GFP positive HIV infected cells were enumerated by fluorescence microscopy using the high content CYTELL imaging platform (GE Healthcare). An average of 25,000 cells were analysed per condition by acquiring multiple fields of view. Under the above conditions, viral transfer is primarily observed to be cell-cell, as supernatants from infected cells cultured alone over this period did not result in significant infection when added to the same target cell types. In addition, the inclusion of the antiretroviral reverse transcriptase inhibitor Efavirenz to co-cultures block eGFP expression within the target cell population (ie. The GFP signal is not consistent with simple endocytosis in the target cell type).

### Quantitative analysis of live cells during viral transfer

#### Cell-cell contacts

The cumulative number of contacts between each infected donor cell and any uninfected target cells (TZM-bl) was determined via frame-by-frame inspection of 3h time-lapse movies. Each contact was recorded as a finished track using ImageJ’s MTrackJ plugin [71]. N>50 cells from two independent experiments were analysed for each group. Cells that exit the field of view were excluded from analysis. Whilst this enumerated absolute number of contacts, cell to cell engagement was enumerated when infected donor cells were contacting the same uninfected target cells for a duration greater than 5 minutes.

#### Cellular circularity vs Gag-polarisation

At the virological synapse, donor-cell circularity was calculated using the circularity measurement function in ImageJ (https://imagej.nih.gov/ij/plugins/circularity.html), where c=1 indicates a perfect circular shape. Gag polarization was calculated as (IntP/IntD)-1, where IntP is the GFP intensity signal at the proximal cell quarter (i.e. contacting the target cell), whereas IntD is the intensity at the distal cell quarter. A polarization value of 1 indicates 100% enrichment in the proximal quarter.

### Immunofluorescence microscopy

For live and fixed cell imaging, 0.5 × 10^6^ cells were infected with HIV stocks at an MOI of 0.1 for 48 hours under standard culture conditions before analysis.

Live cell imaging was performed as described previously using an inverted Olympus IX-70 microscope with 60 × 1.42 NA oil immersion lens and an Evolve 512 back-thinned EM-CCD camera (512*512) (DeltaVision ELITE Image Restoration Microscope, GE Healthcare). For time-lapse movies, approximately 5 × 10^4^ cells were seeded onto a 35mm imaging dish with ibidi polymer coverslip bottom (#1.5) (Ibidi, Martinsried, Germany) and eGFP and DIC channels imaged at approximate 2 frames/sec, with time lapse movies presented as overlays. Manual single particle tracking was done using ImageJ with MTrackJ plugin and filopodial lengths calculated as described previously[9]

For counting virus particles, wells of a 96 well Sensoplate, glass bottom, black (Greiner Bio-One International GmbH) were coated with poly-L-lysine according to manufacturer’s instructions (Sigma-Aldrich). 50µl of virus supernatant was serially diluted at 1 in 5 dilutions steps and then added to each well and plate spun at 2500 x g for 40 minutes at RT followed by fixation with 4% formaldehyde (v/v) for 20 minutes at room temperature. Fluorescent virus particles were then imaged using DeltaVision Elite microscope described above and a total of five fields of view per well were acquired in the GFP channel. Viral particles were then enumerated using ImageJ using the 2D/3D particle tracker in MosaicSuite (MOSAIC Group, Dresden).

For fixed cell imaging, 5 × 10^5^ cells were cytospun onto 22 × 60 mm #1.5 coverslips (VWR international, Batavia, IL), pre-coated with CellTak (Corning), and then fixed in 4% formaldehyde solution (v/v) for 20 minutes at room temperature followed by neutralization with 50 mM NH4Cl (Sigma) for 3 minutes. Cells were permeabilized with 0.05% Triton-X (Sigma) for 1 minute at room temperature, stained with indicated mAbs in the presence of 5% serum, followed by the appropriate secondary antibody. For staining filamentous actin (F-actin), cells were stained with directly conjugated Alexa Fluor-647 Phalloidin (Thermofisher Scientific) for 20 minutes at room temperature prior to mounting in Prolong Gold antifade mountant with DAPI (Thermofisher Scientific). Cells were visualised with a 100 × 1.4 NA oil immersion lens using a DeltaVision Elite microscope and a Photometrics CoolSnap QE camera. Images were acquired as 50 to 60 optical sections, 0.15µm to 0.20µm in thickness, deconvoluted and volume projections generated using DeltaVision SoftWorx software, version 7.0. Unless otherwise stated, all fixed cell data presented herein are Z-series volume projections.

### Correlative FIB-SEM

In order to perform Correlative FIB-SEM, 0.5 × 10^6^ U937 cells were infected with HIViGFP^-Env^ at an MOI of 0.1. 48 hours post-infection, cells were washed twice, resuspended in warm RF-10 and incubated at 37°C, 5% CO2 for a further 30 minutes. This was done to allow the cells to recover the filopodial activity post washing steps. 100μl of cell suspension was then added to a gridded 35mm glass bottom dish (# 2) from MatTek Corporation (MA, USA), that had been pre-coated with CellTak according to manufacturer’s instructions. Cells were cytospun at 80 x g for 1 minute before fixing in fresh 4% (v/v) formaldehyde. Cells were then imaged using DeltaVision Elite microscope, infected cells identified by GFP expression and the grid coordinates noted. Cells grown and imaged on gridded coverslips as described above were further fixed in Karnovsky’s fixative prior to transport. The samples were then post-fixed, stained and resin embedded as previously described [20].

Data collection was performed on a Zeiss Crossbeam 540 FIB-SEM (Zeiss Inc., Germany) controlled by ATLAS3D (Fibics Inc, Canada), as follows. Since the cells were adhered, after the cover slips were removed, the outlines of the cell membranes were just under the top resin surface and could be visualized using a scanning electron beam operated at 3 kV. The pattern of the gridded coverslip was also transferred to the resin surface, allowing correlation with the light microscopic images and location of specific cells of interest. These cells were protected by depositing a layer of platinum on top of the resin, effected by the FIB operated at 700 pA and 30 kV. Notches were FIB milled (at 50 pA) into the platinum to monitor and adjust for specimen drift x, y and z directions as well as beam tuning in real time during data acquisition. Image contrast was achieved by FIB-depositing (at 700 pA) a layer of carbon atop the platinum, and the cells were exposed from the side by FIB milling a trench in the resin at 30 nA followed by 3 nA until the plasma membrane was reached.

During data collection, the FIB and SEM were operated simultaneously, at 30 kV, 700 pA and 1.5 kV, 1 nA, respectively. The EsB detector was operated at a 900V grid voltage to produce a 2-D stack of images composed of predominantly back-scatter electronic signal. The SEM pixel size was set at 3 or 4 nm and total dwell time of 4 µs, and FIB step size set at 9 or 12 nm. The resulting stack of images were aligned, binned, denoised and inverted with IMOD based scripts {Kremer,, 8742726} to yield isotropic 3-D image volumes. Segmentation of these reconstructions was completed using Amira (ThermoFisher Inc) or 3DSlicer {Fedorov,, 22770690}. Sub-volumes of veils and filopodia were extracted from these data and fiducials were placed at the locations of budding virions, which could be easily observed. Thus the number and location of individual virions as well as the surface area of veils and filopodia could be quantified using available modules in the XimagePAQ extension for Amira. This extension was also used to calculate mean curvature of veils and filopodia; areas with negative curvatures were named “base” positive curvature as “ridge” or “tip”, respectively, and neutral curvature as “face” or “shaft”, respectively. Representative features were false colored on a spectrum corresponding to negative (red) to positive (blue) mean curvatures.

### Mass spectrometry of purified virions

Parental and mutant U937 cell lines were infected with HIViGFP^ENV-ve^ at an MOI of 0.3 and after overnight culture cells were washed three time in fresh media to remove the residual inocula. Culture supernatants were then pooled after a harvest at 48 hours and 96 hours post infections. Virion preparations were depleted with antiCD45 paramagnetic microbeads (Miltenyi Biotech cat# 130-045-801) as described previously[32] with following modifications: Clarified cell culture supernatants were incubated with continuous mixing for 1hour at room temperature with anti-CD45 microbeads at a concentration of 4µl beads per ml of supernatant. CD45 immunoaffinity depleted supernatants were then separated from beads by placing in magnetic separators followed by centrifugation at 28,000 x g for 90 minutes at 4°C to pellet virions. Pelleted virions were resuspended in PBS and lysed using 4xLDS sample buffer (BOLT, Invitrogen). CD45 microvesicle depletion was determined by western blotting as previously described [32]. Samples were subsequently separated by SDS-Polyacrylamide gel electrophoresis using 1mm thick 4-12% Bis-Tris gel (Invitrogen), followed by staining with Instant Blue Coomassie stain (Expedeon Inc, USA). Each virion containing lane was divided into 18 contiguous sections and each section then subjected individually to In-Gel digestion protocol{8962070}. Briefly, gel slices were de-stained (50% (v/v) acetonitrile, 50mM NH4HCO3), reduced (10mM DTT in 50mM NH4HCO3) and alkylated (55mM Iodoacetamide in 50mM NH4HCO3) before being trypsinised with 100ng of Trypsin (Promega) for 16h at 37°C. Gel slices were then treated with the following solutions sequentially for 1hour each at RT: 50% (v/v) acetonitrile/0.1% (v/v) formic acid and 100% acetonitrile. Samples were then dried in a centrifugal concentrator before resuspending in 20µl of 0.1% (v/v) formic acid.

#### Mass spectrometry

Proteolytic peptide samples were separated by nano-LC using an UltiMate 3000 HPLC and autosampler system (Dionex, Amsterdam, Netherlands) and eluting peptides ionized using positive ion mode nano-ESI following experimental procedures described previously {22083589}. MS and MS/MS were performed using a Q Exactive Plus mass spectrometer (Thermo Scientific, Bremen, Germany). Survey scans *m/z* 300–1750 (MS AGC = 3×10^6^) were recorded in the Orbitrap (resolution = 70,000 at *m*/*z* 200). The instrument was set to operate in DDA mode, and up to the 12 most abundant ions with charge states of >+2 were sequentially isolated and fragmented via HCD using the following parameters: normalized energy 30, resolution = 17,500, maximum injection time = 125 ms, and MSn AGC = 1×10^5^. Dynamic exclusion was enabled (exclusion duration = 30 s).

#### Sequence database searches and protein quantification

Sequence database searches were performed using MaxQuant (version 1.5.8.0), run using standard parametersusing the Andromeda sequence database search utility. Andromeda was employed using the following parameters: precursor ion and peptide fragment mass tolerances were ±4.5 ppm and ±0.5 Da respectively; carbamidomethyl (C) was included as a fixed modification; oxidation (M) and N-terminal protein acetylation were included as variable modifications; enzyme specificity was trypsin with up to two missed cleavages; and Human sequences in the Swiss-Prot database (February 2017 release; 20,168 Human sequence entries) were searched. Searches were performed with the “match between runs” feature selected, and proteins identified in each gel lane were quantified using the MaxLFQ algorithm employed using standard parameters.

#### Statistical tests

OriginPro (Version 9.0; Originlab Corporation) was used to perform statistical analyses and generate graphs, unless otherwise specified. Unless otherwise stated, the data from 2 groups were compared, normal distributions were tested using a Shapiro□Wilk test and for normally distributed data the probability that the mean of each group was significantly different was evaluated using an unpaired Student’s *t* test. For data that is not normally distributed, the probabilities were determined using the Mann□Whitney *U* test. Unless otherwise stated, statistics are summarised in figures as *p<0.01, **p<0.001 & ***<p<0.0001. All data presented herein is representative of a minimum of 3 independent replicates. Where appropriate, the distribution of raw data is presented and no outlying datasets have been excluded for statistical analysis.

#### Availability of reagents

With the exception of plasmid constructs that encode for full length HIV, the majority of constructs herein will be submitted to Addgene for distribution. Constructs not available on Addgene, will be available upon request. Plasmids encoding full length HIV will be made available upon request, providing the requesting laboratory has the appropriate import permit and biological containment facilities and approvals.

#### Human ethics statement

Herein all authors state and confirm that they have complied with all relevant human ethical regulations outlined under ethics approval # #HREC/13/SVH/145 (St Vincent’s Hospital Sydney, NSW, Australia) to obtain whole blood for research purposes.

#### Data availability statement

Due to the large size of files datasets generated in figures 1, 2, 4, 5 and 7, the data will be available using the Dryad Data Repository.

## References

### Supplementary Figure Legends

**Figure S1. Filopodial network disruption as a result of formin and Arp2/3 depletion**.

Number of filopodia per cell were enumerated using live cell microscopy. In the left panel formins Diaph1, Diaph2 and FMNL1 are depleted using shRNA. Due to lack of detectable expression, other addition known formins were not tested. In the right panel, following depletion of Diaph2, cells were rescued with C/A Diaph2 or co-shRNA depleted (in addition to Diaph2 depletion) for the formins Diaph1, FMNL1 or the Arp2/3 complex. Note, only Arp2/3 + Diaph2 co-depletion decreases filopodial frequency. B. Filopodial lengths derived from the same cells observed in A. Note only Diaph2 depletion and Diaph2 & Arp2/3 co-depletion have significant impacts on filopodial lengths. Also note C/A Diaph2 can rescue filopodial lengths post Diaph2 depletion. Data represents pooled datasets from three independent experiments.***p<0.0001.

**Figure S2. Filopodial frequencies and lengths relative to HIV infection**

A. % Frequency of cells with filopodia. Infected (green) uninfected (grey). Each data point represents a single independent experiment B. Filopodial lengths in the presence of HIV infection. Accumulative filopodial lengths pooled from the 4 independent experiments outlined in A. *p<0.01, **p<0.001

**Figure S3. Analysis of clonal populations for shRNA knockdown of targets**.

A-E Western blotting of cellular lysates using antigen specific antibodies to A. Rac1, B. IQGAP1, C. Arp2, D. Diaph2. In each case, GAPDH immunoblotting of the same transferred lysate is presented below each upper panel. F. Knockdown relative to untreated controls. G. Residual protein expression calculated by determining the band intensities using GeneTools software from Syngene followed by background correction and lane normalization of GAPDH. Remaining protein was then calculated by dividing the intensities of target bands by the lane normalization factor ^17^. H. Metabolic activity (viability) of the main clonal cell lines used within this study. I. Relative Gag-iGFP expression within each clonal cell line. All cells were infected as outlined within quantitative cell to cell virus transfer assays. Cells were fixed and then mean fluorescence intensity quantified by flow cytometry. Expression of Gag-iGFP is then expressed as % relative to the WT cell line control. In H and I, data is representative of a minimum of three independent control experiments.

**Figure S4. Verification of CRISPR/Cas9 gene-editing by DNA sequencing**. U937 cells expressing Cas9 and a specific gRNA where clonally expanded, and clones were screened by genomic PCR amplification of the gRNA target site followed by DNA sequencing. Shown are representative examples for different gRNAs. In each case, the wildtype genome is shown above the mutated genome for comparison. The 20 bp region of the gRNA complementary to the host wildtype genome (protospacer) is highlighted in orange, whereas the protospacer adjacent motif (PAM sequence) is indicated by a green box (NGG in lead strand or CCN in complementary strand). **A**. Wave2^**k/o**^ cell clone with three point mutations and 1 bp deletion in positions 14-17 of the protospacer (red box). Single chromatogram peaks throughout the amplicon indicate homozygous mutant alleles. The 1b deletion leads to a frameshift from residue 74, resulting in >70% loss of native protein sequence and a premature stop codon at residue 127 of 498. **B**. Cdc42^**k/o**^ cell clone with heterozygous mutations, including at least 1 bp deletion per allele at positions 17-20 of the protospacer. Frameshift at these positions leads to >80% loss of native protein sequence.

### Supplementary tables

**Table S-I. Statistical Summary for Unpaired two tail T-Test for HIV bud frequency relative to cortical F-Actin structures**

**Table S-II. Plasmid constructs generated and used in this study**

**Table S-III. shRNA sequences used in this study**

**Table S-IV. CRISPR gRNA used in this study**

**Table S-V: List of selected proteins detected in virion preparations from Diaph2**^**+ve**^**Arp**^**+ve**^ **Diaph2**^**-ve**^**Arp**^**+ve**^ **cells and Diaph2**^**-ve**^**Arp**^**-ve**^ **cells**.

**Table S-VI: Fold change in the levels of cellular proteins detected in virion preparations from Diaph2**^**-ve**^**Arp**^**+ve**^ **cells and Diaph2**^**-ve**^**Arp**^**-ve**^ **cells relative to Diaph2**^**+ve**^**Arp**^**+ve**^ **cells**.

### Supplementary movie files

**Movie 1. FIB-SEM imaging reveals extensive short filopodial networks in the absence of Diaph2**.

Rotation of 3 dimensional rendering from accumulative *FIB-SEM* datasets derived from cells depleted of Diaph2. At the conclusion of the rotation, filopodial networks are highlighted individually in colour. Note the extensive curvature and sub-populations of short filopodia that have branching (i.e. Consistent with the action of Arp2/3).

**Movie 2. FIB-SEM imaging reveals extensive lamellipodial networks in cells with co-depleted Diaph2 and Arp2/3**.

Rotation of 3 dimensional rendering from accumulative *FIB-SEM* datasets derived from cells shRNA co-depleted of Diaph2 and Arp2/3. At the conclusion of the rotation, lamellipodial networks are highlighted individually in colour. For this dataset, we have used a HIV infected sample to highlight the enrichment of virions to the ridges of lamellipodial network (see timestamp 14s and the ridge of the green highlighted). Of note, HIV infection does not influence cortical F-Actin in this cellular clone and this image is representative of HIV negative and positive samples.

**Movie 3. Diaph**^**C/A**^**filopodia are straight, dynamic filopodia that exclude HIV**.

Live imaging of HIViGFP^+ve^ (green) Diaph^C/A^-mCherry (red) cells presented as an overlay with Differential Interference Contrast imaging to capture filopodial networks. Frame rate is presented in real-time at 0.841 frames per second.

**Movie 4. Diaph**^**C/A**^ **filopodia cannot be inhibited by HIV curvature mutants**

Live imaging of Diaph^-ve^ HIViGFP^CA99A+ve^(green) cells (left panel) and Diaph^C/A^-mCherry (red), HIV^CA99A+ve^ (green) cells presented as an overlay with Differential Interference Contrast imaging to capture filopodial networks. Frame rate is presented in real-time as in the figure legend to Movie 3. Note, in the left panel, the complete lack of any filopodial network, whereas in the right panel Diaph^C/A^ can readily induce filopodia in the present of HIVCA99A+ve.

**Movie 5. HIV augmented filopodia in WAVE2**^**k/o**^ **cells tether and then retract upon cell-cell conjugation**. Tethering capacity of augmented filopodia is presented. Herein a thick filopodial structure at approximately 40μm in length is presented and targets cell that subsequently undergoes cell division. Note, that all filopodial activity ceases, once the infected donor cell has conjugated between the two recipient daughter cells. Movie is a time lapse speedup with each frame at 30 seconds intervals.

**Movie 6. Filopodial networks in HIV infected WAVE2**^**k/o**^ **cells collapse prior to cell-cell HIV transfer**. Whilst extensive filopodial activity proceeds upon initial cell-cell contact, at the movie time point 31s, all filopodial activity ceases with retraction of all filopodia. Immediately following filopodial retraction, GFP delivery/release is firstly observed, followed shortly after by cell-cell fusion and syncytia formation. Movie is a time lapse speedup with each frame at 30 seconds intervals.

